# Dissociating the effects of distraction and proactive interference on object memory through tests of novelty preference

**DOI:** 10.1101/2020.08.16.253179

**Authors:** K. Landreth, U. Simanaviciute, J. Fletcher, B. Grayson, R.A. Grant, M.H. Harte, J. Gigg

## Abstract

Encoding information into memory is sensitive to distraction whilst retrieving that memory may be compromised by proactive interference from pre-existing memories. These two debilitating effects are common in neuropsychiatric conditions but modelling them preclinically to date is slow as it requires prolonged operant training. A step change would be the validation of functionally equivalent but fast, simple, high-throughput tasks based on spontaneous behaviour. Here, we show that spontaneous object preference testing meets these requirements in the subchronic phencyclidine (scPCP) rat model for cognitive impairments associated with schizophrenia. scPCP rats show clear memory sensitivity to distraction in the standard novel object recognition task (stNOR). However, due to this, stNOR cannot assess proactive interference. Therefore, we compared scPCP performance in stNOR to that using the continuous NOR task (conNOR), which offers minimal distraction, allowing disease-relevant memory deficits to be assessed directly. We first determined that scPCP treatment did not affect whisker movements during object exploration. scPCP rats exhibited the expected distraction stNOR effect but had intact performance on the first conNOR trial, effectively dissociating distraction by using two NOR task variants. In remaining conNOR trials, scPCP rats performed above chance throughout but, importantly, their detection of object novelty was increasingly impaired relative to controls. We attribute this effect to the accumulation of proactive interference. This is the first demonstration that increased sensitivity to distraction and proactive interference, both key cognitive impairments in schizophrenia, can be dissociated in the scPCP rat using two variants of the same fast, simple, spontaneous object memory paradigm.

## Introduction

The encoding of new information into memory depends on being able to stay ‘on track’, that is, to use focused attention to resist extraneous distraction. Retrieval of successfully encoded new memories then requires the ability to prevent proactive interference, that is, contagion from other, similar memories, disrupting current memory representations and decision making (Postle and Brush, 2004; Postle et al., 2004). Sensitivity to both of these factors is a core symptom of several neuropsychiatric disease, including early Alzheimer’s disease (Bondi et al., 1994). Indeed, cognitive impairments associated with schizophrenia (CIAS) include cognitive control (Nuechterlein et al., 2004) and its core executive function of inhibitory control, which includes focussed attention (resistance to distraction) and cognitive inhibition (Diamond, 2013). Impaired resistance to proactive interference may be an underlying factor for a variety of memory impairments in schizophrenia (Park et al., 2003; Heckers et al., 2000; Burton et al., 2018; Ettinger et al., 2018; Girard et al., 2018). These symptoms are severely debilitating (Elvevag and Goldberg, 2000; Green, 1996), occur prior to psychosis (Kahn and Keefe, 2013) and are resistant to treatment (Keefe et al., 2007; Nuechterlein et al., 2004). The CNTRICS initiative identified attention and cognitive inhibition as symptom areas with high translational potential for preclinical research (Barch et al., 2009) and several preclinical, operant-based tasks with translational validity have been developed to meet this need (Gilmour et al., 2013). However, whilst these tasks highlight rule-based, attentional and interference features, they also commonly require weeks to months of training to attain threshold performance. Ideally, our preclinical task armoury to investigate distraction and interference effects should also include high-throughput, simple, short-duration tasks that require only spontaneous behaviour.

The novel object recognition (NOR) paradigm has been used most commonly to characterise the subchronic phencyclidine (scPCP) model of CIAS (Cadinu et al. 2018; Neill et al., 2010; Neill et al., 2014; Lisman et al., 2008). The standard NOR task (stNOR) depends on maintaining object information in short-term memory and scPCP treatment induces a NOR deficit in mice (Nagai et al., 2009; Hashimoto et al., 2008; Gigg et al., 2020) and rats (Horiguchi and Meltzer, 2012; Snigdha et al., 2010) that is produced by distraction during the inter-trial interval (Grayson et al., 2014; Gigg et al., 2020). This supports the conclusion that, whilst the scPCP model can encode object memory, maintaining this information is abnormally sensitive to disruption. Whilst this demonstrates the cross-species validation of a simple and fast means to measure distraction susceptibility, this effectively precludes stNOR as a probe for disease-relevant memory deficits. However, recent developments allow NOR to be probed in a continuous trials design (conNOR); allowing the sequential performance of 10 or more object preference trials in a single session, all with minimal distraction between task phases (Albasser et al., 2010; Ameen-Ali et al., 2012; Chan et al., 2018). Using such an approach provides a promising means to probe object memory deficits independent of distraction in the scPCP and other models of human neuropsychiatric conditions.

Here, we first ensured that scPCP treatment had no *a priori* effect on object sensing through whisker movements. This was important as whisker kinematics provide key tactile information to rodents about the environment and are altered in other rodent disease models (Grant et al., 2020; Grant et al., 2014; Simanaviciute et al., 2018). We next ran stNOR to reproduce the effect of distraction in scPCP rats. Following this, rats experienced eleven trials of a single conNOR session to test the effect of proactive interference on memory performance without the potential confound of distraction. Our hypothesis was that we could achieve behavioural dissociation for the effects of distraction and proactive interference on object memory by comparing scPCP performance between these two related, simple, spontaneous tasks. A positive result here would add further relevance to the scPCP model for pre-clinical research and provide attractively simple and high-throughput methods to probe high-level cognitive deficits relevant to schizophrenia.

## Methods

### Animals

Experiments were performed using 20 female Lister hooded rats (Charles River, UK; 190-224g at study start). Animals were housed in groups of five in standard housing conditions (Techniplast ventilated cages, temperature 20°±2°C and humidity 55±5%, University of Manchester Biological Services Facility) with *ad libitum* access to standard chow and water. All experimental procedures were carried out in the light phase of their cycle (09:00-14:00) and performed under Home Office UK project licence in accordance with the Animals (Scientific Procedures) Act UK 1986 and approved by the University of Manchester AWERB (Animal Welfare and Ethical Review Body). A summary of all experimental stages is provided in Figure 1A.

**Figure 1.**
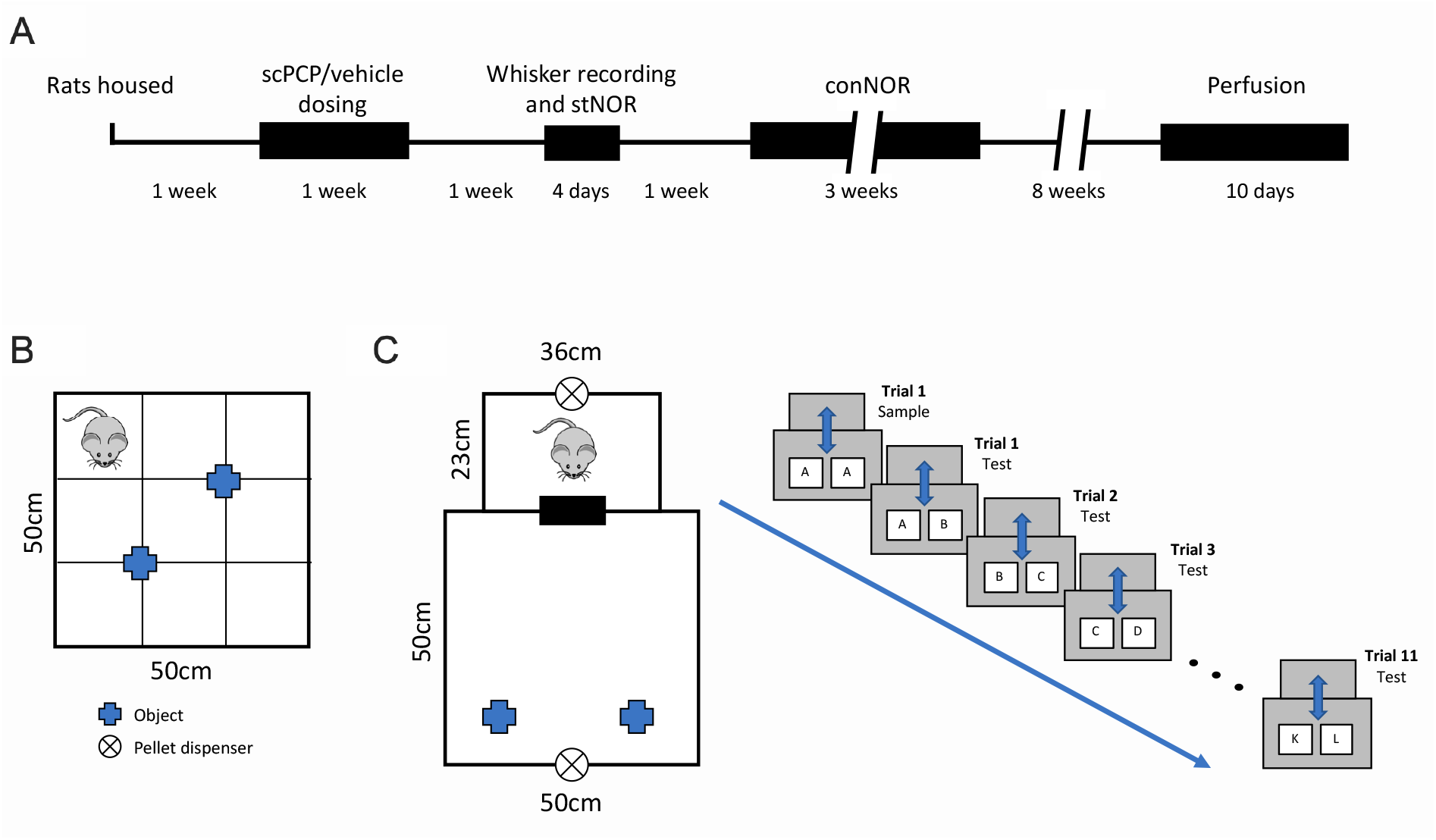
Experimental design. A. Timeline for the study. B. The stNOR arena. C. The arena (left) and the trial design (right) for conNOR testing. Different pairs at test are displayed as A+A, A+B, etc.

### Dosing

10 rats were dosed with phencyclidine hydrochloride (2mg/kg, i.p.) and the other 10 with vehicle (0.9% saline, i.p.). Dosing for all rats followed a standard sub-chronic regimen with injections delivered twice daily for seven days, followed by a seven-day washout period (Grayson et al, 2014).

### Whisker movements

Whisker movements were filmed after dosing to determine whether scPCP treatment changed the way the rats explored objects. Testing was carried out as described previously (Grant et al., 2014; Simanaviciute et al., 2018). Rats explored two different objects sequentially (5min each) for one 10min session in a 30 × 50 × 15cm transparent Perspex enclosure sitting on a lightbox (59.4 × 84.1cm). Objects (14.3 × 6 × 3cm) were composed of three smooth plastic toy bricks with 50% of these covered in tape to provide a different texture. The box and objects were cleaned with 70% ethanol between rats. Whisker movements were tracked at 500 frames per second by an overhead high-speed video camera (Phantom Miro ex2, resolution 640×480 pixels). Video clips were collected for each rat by manual trigger when it moved towards an object. Clips were trimmed and included for analyses if they fit published criteria where: i) the rat was clearly in frame; ii) whiskers on both sides of the face were visible; iii) the head was parallel to the floor (no extreme pitch or yaw, no object climbing); iv) there were at least 50 frames of contact with the object (no contact with the walls of the box); v) the tracked portion of the clip was at least 150 frames long (Grant et al., 2014). Since this investigation was only concerned with object exploration, an additional criterion was added; i) the rat was contacting the object with their whiskers, with no additional whisker contact on the walls of the box. After clip selection, 1-8 clips/rat, 149-1363 frames/clip, 0.298-2.726 seconds/clip were analysed using the Automated Rodent Tracker v2 (Gillespie et al., 2019). Image analysis located the tip of the rodent’s nose and the body centroid (Figure 2). A co-registered scale bar provided a calibrated measure of locomotion speed. For whisker tracking, the software automatically found the snout orientation and position and the whisker angles relative to cranial midline (Figure 2). Whisker angles were measured as the angle between whisker and mid-line of the nose and head; larger angles represented more forward-positioned whiskers. Tracking was validated by manual inspection of video footage. Clips were removed where the tracked portion of the clip was less than 100 frames long, leaving 233 clips (106 vehicle and 127 scPCP, 119 smooth and 114 textured, 112 before and 121 after injections) for analysis.

Mean whisker angle was calculated as a frame-by-frame average of all tracked whiskers on each side of the face. The mean whisker angles allowed the tracker to calculate the following whisker position and movement parameters: mean angular position (the average whisker angle, measured in degrees), amplitude (2√2* the standard deviation of whisker angles, to approximate the range of whisker movements in degrees), asymmetry (the difference in the mean angular position between all whiskers on left and right sides; degrees), and the mean angular retraction and protraction speeds (calculated as the average velocity of all the backward (negative) and forward (positive) whisker movements, respectively; degrees/s) (Gillespie et al., 2019). For mean angular position, amplitude, and retraction and protraction speeds, the values for right and left whisker measurements were averaged.

## Object memory assessments

### Standard novel object recognition memory test (stNOR)

After the 7-day dosing washout period animals were subjected to a single novel object recognition trial (Ennaceur and Delacour, 1988) to ensure that scPCP rats expressed the expected NOR deficit (Grayson et al., 2007). The NOR arena was a 50cm^3^ box marked into a 3×3 grid on the floor (Figure 1B). A separate holding box was positioned close to the apparatus, consisting of a 15cm×20cm opaque plastic tub into which animals were placed during the inter-trial interval. Rats were first habituated to the test arena by placing them individually into the box for 15min. The next day, rats were subject to one stNOR session consisting of a 3min object acquisition phase (explore two identical copies of a novel object), removal to the holding box for a 1min inter-trial interval (ITI) then return to the test arena for a final 3min test phase (explore a copy of the acquisition object and one novel object). Objects were well validated for equal baseline preference in prior testing (Grayson et al., 2007) and placed at positions equidistant from where the rat was introduced to the arena to ensure no spatial bias. Exploration was recorded via overhead video camera and analysed offline using the Novel Object Recognition Task Timer (https://jackrrivers.com/program/). Rats were considered to be exploring when the nose was pointed towards and within 2cm of an object; however, any time spent climbing on the object was discounted. The discrimination index (DI) between novel and familiar objects at test (stNOR) was measured by calculating the difference between exploration time for the two objects at test and dividing the result by the sum of their exploration time. Therefore, the more positive the value, the more exploration of the novel object at test, with negative values representing enhanced exploration of the familiar object.

### Continuous novel object recognition memory test (conNOR)

The conNOR apparatus consisted of a holding chamber attached to a larger experimental chamber by a computer-controlled sliding door (Campden Instruments Ltd., UK). An overhead video camera recorded exploration within the experimental chamber. Computer-controlled pellet dispensers with reward troughs were placed in both chambers on the walls furthest from the door (Fig 1C) to deliver standard rodent tablets (LabTab AIN-76A, 45mg; TestDiet, USA). Task sequences and operation of the door, pellet dispensers and recording of behavioural video and visits to retrieve pellets were managed and recorded by ABET II software (Campden Instruments Ltd. UK). The advantage of this arena for present purposes is that it does not require any handing during the ITI; animals are only handled twice, once at the start and the for final removal once all trials have finished.

Rats were first habituated to the conNOR arena over four days, during which timed dispensing of food pellets encouraged shuttling between holding and experimental chambers (Chan et al., 2018). On day 1, cage groups were placed into the arena with the central door open for 30min to explore freely, encouraged by single pellet dispensing into both chambers every minute. On day 2, rats were placed singly into the holding chamber with the gate open for 20min of free exploration with pellets delivered to both chambers every 1min. Animals that fed in and explored both chambers moved to habituation day 3 where they were placed in the holding chamber, a pellet was delivered and, when taken from the dispenser, the door opened and a further pellet was delivered in the experimental chamber. Once the animal shuttled successfully the door closed, opening again once the pellet had been taken from the experimental dispenser. This process continued for 20min, with animals moving on to habituation day 4 if they shuttled quickly between chambers at least 18 times. On habituation day 4 rats were placed individually into the holding chamber and two identical objects were placed in the experimental chamber. After taking a pellet and shuttling to the experimental chamber the rat could explore these objects for 5min after which the door re-opened and a pellet was delivered to the holding chamber. Once the rat returned to the holding chamber for a 1min ITI, the door closed and objects in the experimental chamber were exchanged for a second pair of novel objects (if the animal failed to re-enter the holding chamber within 3min it was moved back to the habituation day 3 protocol). This protocol was repeated for a third pair of objects, with animals being required to actively explore the objects to be permitted to move on to the testing phase.

Rats were mildly food restricted on the evening prior to testing (12g/rat/day standard chow) to encourage performance in the conNOR arena the next day. Animals were placed in the holding chamber and a pellet was delivered. Once the pellet was taken from the dispenser the door opened and 1min later a pellet was dispensed in the experimental chamber (apart from taking a pellet to initiate the testing sequence they were not required to retrieve or consume pellets at any other point during testing). The door closed once the rat had shuttled to the experimental chamber, which now contained two identical novel objects (A+A). Objects in all trials were all of a similar in size and constructed of plastic, glass or ceramic. Rats explored the A objects for 2min before the door re-opened and a pellet was delivered to the holding chamber, prompting the animal to shuttle and the door to close. During a 1min ITI the objects were removed and replaced with an identical object A and a novel object B. At the end of the ITI the door re-opened and a pellet was delivered to the experimental chamber, permitting a further 2min object exploration period. This process of shuttling between holding and experimental chambers was repeated over 11 object pairs (A+A, A+B, B+C … K+L; Fig 1C) to allow for continuous testing of object memory. The side of the chamber on which the novel object was placed was counterbalanced within rats, along with the object order between rats. Object exploration was measured from recorded video as per stNOR. For analyses the DI metric was again used for each trial where exploration times were summed for all trials to that point. For example, for conNOR trial 3 the sum of exploration durations for familiar objects on trials 1 to 3 was subtracted from the sum of novel object exploration over the same trials, the result divided by the sum of these two measures (Chan et al., 2018).

### Tissue collection and immunohistochemistry

Rats were anaesthetised with isoflurane and then perfused transcardially with phosphate buffered saline (PBS). The brains were collected, cut in the coronal plane to provide blocks that included mPFC (prelimbic and infralimbic cortices) and post-fixed in 4% formaldehyde for 72 hours at 4°C. Once fixed, brains were immersed in 30% sucrose until they sunk and then stored at −80°C until sectioning. Sections were cut at 30μm using a cryostat (Leica Biosystems). One in four serial sections of the prefrontal cortex were suspended in cryoprotectant (30% ethylene glycol, 30% glycerol, 10% PBS, 30% dH2O) and stored (−20°C) until stained. Sections were washed three times in PBS for five minutes each and then transferred into a hydrogen peroxide solution for 30min (0.6% H_2_O_2_, 0.1% triton x-100, 8.8% PBS, 10% methanol, 80.5% dH_2_O). The sections then underwent PBS wash for 5 minutes followed by protein block for one hour (5% normal horse serum, 0.4% triton x-100, 94.6% PBS). Incubation was then started with parvalbumin primary antibody (1:5000; Swant, Switzerland) diluted in protein block solution at 4°C for 36hr.

After incubation, samples were washed twice in PBS and incubated for 2hr with secondary antibody (biotinylated anti-mouse; 1:200; Vector Laboratories; in protein block solution) and washed again in PBS. Sections were then incubated with a Vectoastain ABC kit (Vector Laboratories, UK), in the dark, for 2hr. After a final PBS wash, samples were visualised using DAB substrate kit (Vector Laboratories, UK). Samples were incubated for up to 15 minutes and transferred into distilled water to stop DAB staining. Sections were mounted onto slides and left to dry overnight. Samples were dehydrated using increasing concentrations of ethanol (70%, 90% and 100%) followed by Histoclear (5min) and allowed to dry for 30 minutes then mounted using DPX (Sigma-Aldrich).

Images were viewed on an Olympus BX51 microscope and analysed using ImagePro Plus (v6.3.0.512, MediaCybernetics, USA). For each section the region of interest (PFC) was delineated manually. We then used a 2-d stereological method whereby 35 randomly selected field of views were analysed within this region at 20x magnification using a motorised stage. Within each field a square counting box with inclusion and exclusion lines was used to determine the number of PV positive interneurons (each box was 120×120μm). The data are presented as cell density per mm^2^. All analysis was conducted with the experimenter blinded to treatment condition.

### Data Collection and Statistical Analysis

Videos of whisker movement were tracked and object exploration scored when still blind to treatment condition. All data were analysed by either unpaired two-tailed Students t-test or repeated measures two-way ANOVA with *post hoc* comparisons (Sidak). All analyses were performed using GraphPad Prism (v6.0).

## Results

### Whisker movement assessment

We tested whether scPCP treatment had any effect on whisker movement by filming object encounters for all rats after dosing. Firstly, there was no effect of object texture on our measurements, so data from both object types were combined. As can be seen in Figure 2, for these combined measures we observed no differences in whisker movements between vehicle and scPCP treated rats (two-tailed unpaired t test). Overall, these results show scPCP treatment has no effect on how rats explore objects through active whisker movement.

**Figure 2.**
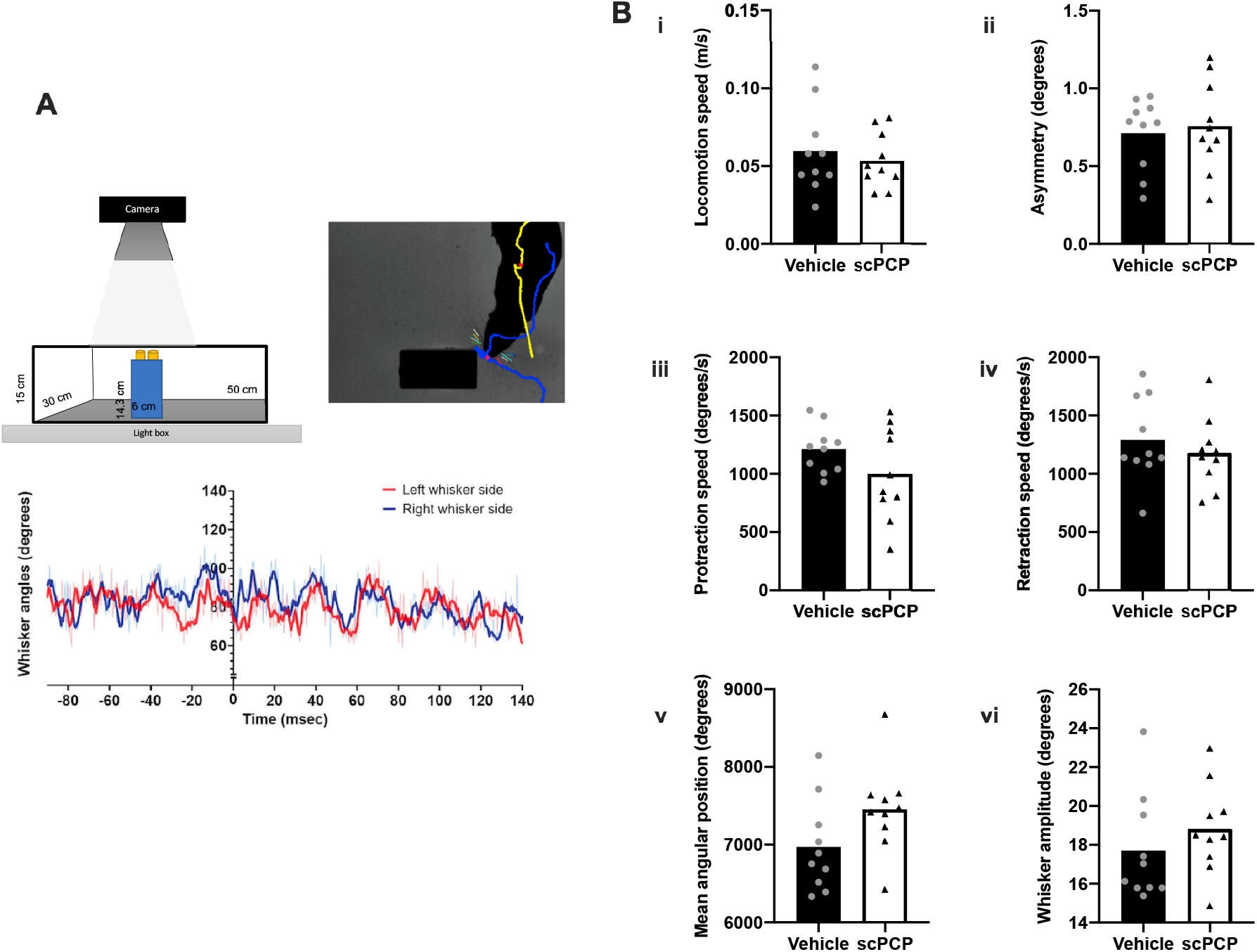
Whisker movement in scPCP and Vehicle rats. A. Top left panel: Diagram of object exploration arena; Top right panel: video frame taken from typical whisker contact with an object to show tracking of whiskers, tip of nose (blue) and body centroid (yellow); Bottom panel: example of whisker tracking;object contact occurred at time x=0; therefore, whisker measurements were extracted from after this time. B. scPCP and vehicle groups were similar in terms of: (i) general locomotion speed in the arena, (ii) asymmetry of whisker position; (iii) whisker protraction and (iv) retraction speed; (v) mean angular whisker position; and (vi) whisker amplitude. N=10 anmals for both vehicle and scPCP groups, with data presented as individual values plotted over group mean.

### Object memory testing using the standard novel object recognition paradigm (stNOR)

During the initial acquisition phase there were no differences in total exploration time between vehicle and scPCP groups; the exploration time for each of the two identical objects at acquisition was also similar both between objects and treatment groups (data not shown). When comparing exploration times of novel versus familiar objects at test (Fig 3A) there was a significant effect of object novelty (F(1,18) = 10.53, p<0.01) and interaction between object novelty and group (F(1,18) = 5.666, p<0.05) with *post hoc* tests revealing significantly higher exploration of novelty in vehicle-treated rats only (p<0.01, Sidak). This result was also seen when comparing exploration as DI preference score (Fig 3B) with vehicle rats showing a significantly higher DI score compared to scPCP (p<0.05, unpaired t test) and only vehicle animals showing DI scores above chance (DI=0; p<0.001, one sample t test). Overall, these results show that scPCP rats displayed the expected stNOR deficit (Grayson et al., 2007).

**Figure 3.**
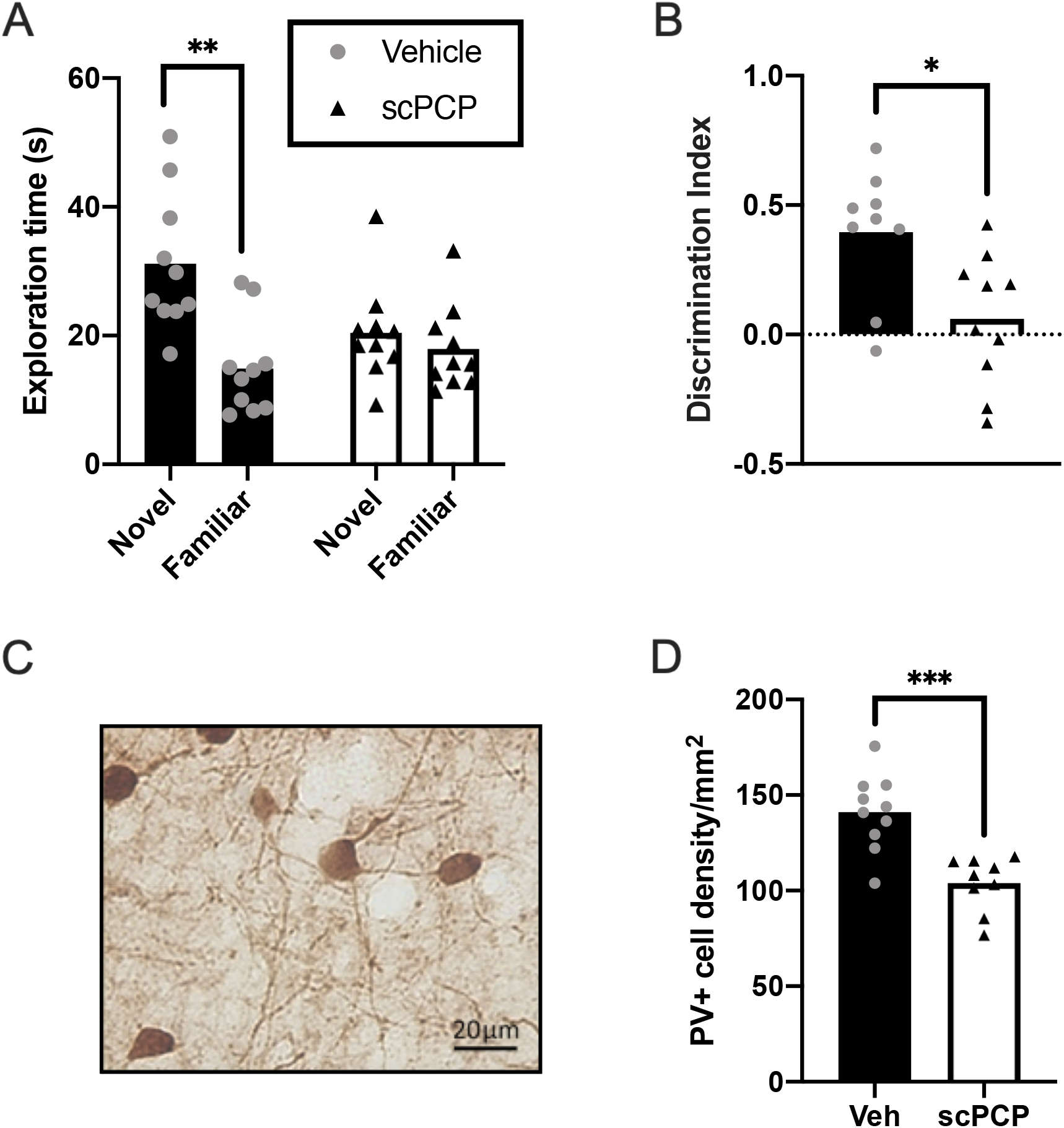
Standard NOR (stNOR) and PV deficit in scPCP rats. A. Exploration times for novel and familiar object at stNOR test; only vehicle rats show a preference for the novel object with scPCP rats exploring both novel and familiar objects similarly. B. stNOR test performance plotted as discrimination index (DI); vehicle rats show significantly higher DI compared to scPCP. Only vehicle group performance was greater than chance. C. Typical example of parvalbumin staining from mPFC in rat. Image x20. D. The density of cells positively stained with anti-parvalbumin antibody is lower in scPCP group. N=10 for both vehicle and scPCP groups with their data presented as individual values plotted over group mean; *p<0.05, **p<0.01, ***p<0.001.

### Object memory testing using a continuous novel object recognition paradigm (conNOR)

Analysis of exploration time over the course of the conNOR protocol showed an effect of trial (F(5.276,94.97)=12.33, p<0.001; Geisser-Greenhouse correction for sphericity epsilon 0.5276) but no effect of treatment and no interaction (Suppl Fig 1A). There was no effect of PCP treatment on mean object pair exploration time (Suppl Fig 1B) and when comparing mean exploration times from the first versus the last four trials (Suppl Fig 1C) there was a decline in exploration time between trial blocks (F(1,6)=17.04, p<0.01) but no effect of treatment or interaction. Thus, although object exploration times tended to decrease and then plateau over conNOR trials, total object exploration for each trial was similar between vehicle and scPCP rats.

The rats’ memory performance for conNOR is seen in the cumulative DI results, which show clearly that both scPCP and vehicle animals were able to recognise novelty throughout all 11 trials (Fig 4A). However, from trial 3 onwards the mean performance of scPCP rats decreased and then plateaued at a lower DI level compared to vehicle. This pattern is supported by a significant effect of trial (F(10,180)=13.48, p<0.001) and interaction between trial and treatment (F(10,180)=2.33, p<0.05) but no *post hoc* differences between scPCP and Vehicle groups for any trial (Sidak). In addition, when considering initial versus late performance by averaging cumulative DI across the first versus the last 4 trial blocks (Fig 4C) there was an effect of trial block (F(1,18) = 43.20, p<0.01) and an interaction between treatment and trial block (F(1,18) = 8.013, p<0.05). Post hoc tests (Sidak) revealed significant differences between these 4-trial blocks for both Vehicle (p<0.05) and scPCP (p<0.001) treatment groups. Thus, whilst both groups showed good object memory throughout conNOR testing there was a significant performance deficit in the scPCP group compared to Vehicle that became more pronounced with increasing numbers of trials (proactive interference).

**Figure 4.**
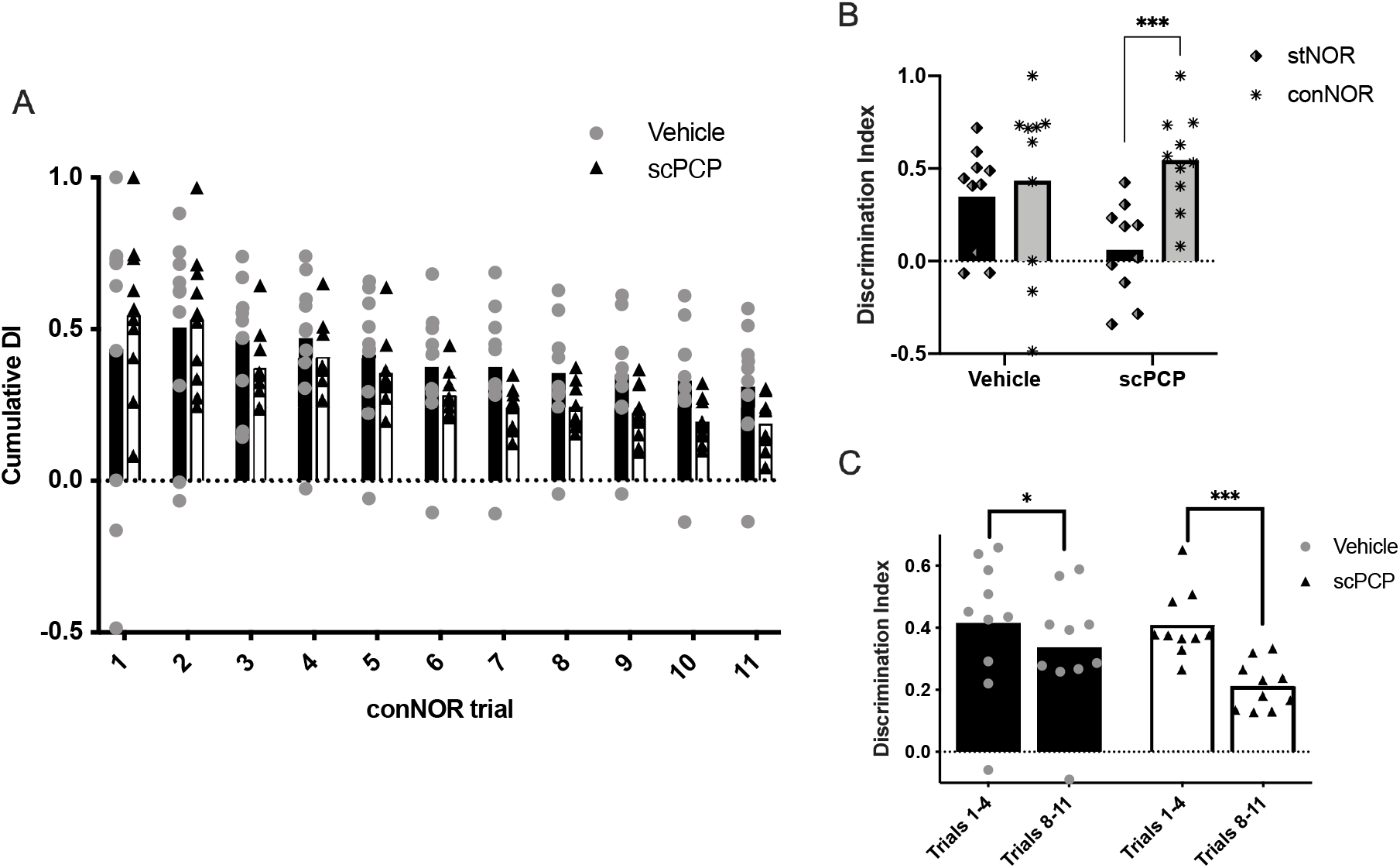
Continuous NOR (conNOR) performance in vehicle and scPCP rats. A. Vehicle and scPCP groups show intact performance throughout the conNOR session but scPCP performance declines from vehicle group from trial 3 followed by a more consistent lower mean value from trial 7. B. The stNOR deficit in scPCP rats and their intact performance on conNOR trial 1 supports the specific effect of distraction in this group for object memory in stNOR but not conNOR. C. Decrease in performance for scPCP rats over trials without distraction in conNOR supports their increased sensitivity to proactive interference compared to vehicle group in conNOR. N=10 for both vehicle and scPCP groups. N=10 for both vehicle and scPCP groups with their data presented as individual values plotted over group mean; *p<0.05, ***p<0.001.

To determine whether we could reproduce the effect of distraction on the persistence of object memory after scPCP treatment we compared performance of stNOR (distraction) to that of conNOR trial 1 (no distraction; Fig 4B). These data show a clear pattern where vehicle rats perform well in both NOR versions whereas scPCP rats show a specific deficit in stNOR only. Thus, there was an effect of the type of NOR test (stNOR vs conNOR; F(1,18)=14.56, p<0.01) and an interaction between NOR test and treatment (F(1,18)=7.155, p<0.05). Post hoc (Sidak) tests showed that there was a significant difference between stNOR and conNOR performance for the scPCP group only (p<0.001). In addition, apart from scPCP rats undergoing stNOR all other groups performed significantly above chance (DI=0; one sample t test; stNOR and conNOR for vehicle both p<0.05, conNOR scPCP p<0.01). These data clearly show that object memory deficits in scPCP rats are sensitive to both distraction and proactive interference and that these effects can be dissociated behaviourally by using two related tests of object preference - the stNOR and conNOR paradigms, respectively.

### Anatomical changes in the scPCP mouse prefrontal cortex

Brains from all animals were analysed for changes in the density of PV-positive cells in PFC (Fig 3C,D). There was a significant reduction for scPCP rats (p<0.001; two-tailed unpaired t test) compared to vehicle, which agrees well with prior validation of the scPCP model.

## Discussion

The acquisition and flexible use of new memory is susceptible to several disruptions of cognitive control. These include difficulty in maintaining attention on new information in the face of distraction and contagion by related, older memories when trying to recall newly encoded information (Postle and Brush, 2004; Postle et al., 2004). These sensitivities to distraction and proactive interference, respectively, are debilitating symptoms in a range of neuropsychiatric diseases; indeed, they are likely to be fundamental factors in underpinning a wide range of cognitive impairments associated with schizophrenia (CIAS)(Girard et al., 2018). CIAS are difficult to treat with current therapies (Nuechterlein et al., 2004; Keefe et al., 2007), so there is a pressing need for new drug discovery in this area of symptomatology. A core pre-requirement for the latter is a range of preclinical models that show CIAS-related deficits in simple, high-throughput tasks that would allow rapid testing of novel compounds whilst maintaining high translational validity. Currently, testing for cognitive control deficits in models of CIAS requires training of rodents in complex operant tasks that often takes weeks to months before animals reach criterion prior to subsequent testing. Whilst these tasks are reliable and offer high translational validity, they are low-throughput. Ideally, our preclinical armoury of tasks to investigate cognitive control aspects such as resistance to distraction and interference would also include high-throughput, simple, short-duration tasks that probe these aspects of cognition by instead relying on spontaneous behaviour. An obvious candidate here is the highly popular novel object recognition paradigm (NOR), which takes advantage of the innate preference of rodents to explore novelty (Ennaceur and Delacour, 1988). Here, we tested whether disease-relevant sensitivity to distraction and proactive interference could be dissociated in variants of the NOR task using a rat model of CIAS. A successful outcome would provide powerful new options for future drug discovery programmes through simple, high-throughput task options, each sensitive to a selective aspect of memory impairment in schizophrenia.

Our chosen model was the sub-chronic phencyclidine (scPCP) treated rat, developed to model the N-methyl-D-aspartate (NMDA) receptor hypofunction hypothesis of schizophrenia (Steinpreis, 1996; Krystal et al., 1994; Javitt and Zukin, 1991; Coyle, 2006). The scPCP rat model effectively produces CIAS-associated neurobiological and cognitive impairments (Neill et al., 2010; Neill et al., 2014; Lisman et al., 2008). Thus, both rat and mouse scPCP models show a robust decrease in parvalbumin (PV) expression (Abdul-Monim et al., 2007; Reynolds and Neill, 2016; Gigg et al., 2020), consistent with inhibitory neurone changes in the cortex and hippocampus of schizophrenic patients (Beasley and Reynolds, 1997; Beasley et al., 2002; Benes et al., 1991). A defining feature of the scPCP model is a deficit in the standard NOR (stNOR) test (Nagai et al., 2009; Hashimoto et al., 2008; Gigg et al., 2020; Horiguchi and Meltzer, 2012; Snigdha et al., 2010). scPCP performance is susceptible to task-related distraction during the stNOR inter-trial delay, supporting the conclusion that initial object memory encoding is good in scPCP rodents but their ability to maintain this information is more sensitive to disruption compared to controls (Grayson et al., 2014; Gigg et al., 2020). Thus, studies using drugs to improve stNOR performance in the scPCP model in particular and other models more generally should take into account the possibility that effects may be via improving attention rather than modulating memory *per se*. Whilst the present data agree with prior studies that scPCP rodents have no deficit in acquiring and maintaining object memoryin the absence of distraction, we tested whether there was a quantitative difference in how scPCP rats use their whiskers to sense this information. Whisker movements during object contact are a vital source of environmental information for rodents and these are abnormal in other rodent models of neuropsychiatric disease that express sensory, motor and cognitive deficits (Grant et al., 2014; Grant et al., 2020; Simanaviciute et al., 2018; Garland et al., 2018). We measured various aspects of whisker movement in vehicle- and PCP-treated rats and found no differences, supporting the view that object-touch related information is preserved at the earliest stages of sensory input. We were confident that that the scPCP phenotype was established in these rats, demonstrated by their inability to discriminate novelty in the single trial NOR test (see below) and decreased density of parvalbumin-positive neurones within the medial PFC. Therefore, we suggest that the lack of whisker movement deficit following scPCP induction here may be due to using an adult rather than developmental model. Indeed, there is ample evidence for sensorimotor acquisition and integration disturbances in schizophrenic patients, potentially affecting the sense of self and body boundaries in particular (Postmes et al., 2014), and many transgenic preclinical models express changes in whisker movements and somatosensory cortical plasticity (Simanaviciute et al., 2018; Greenhill et al., 2015). Although whisker movements can be easily tracked and quantified to measure movement deficits, they are also relatively robust to neural changes (Garland et al., 2018; Grant et al., 2014; Grant et al., 2020), with forms of whisker movements always present, even in late-stage transgenic models. Nevertheless, at least from a behavioural perspective, the acquisition of whisker-related sensory input appears normal in the adult scPCP rat here.

We were next able to reproduce the documented sensitivity of stNOR performance in the scPCP model to distraction (Gigg et al., 2020; Grayson et al., 2014). This was determined by directly comparing stNOR performance to that of the first continuous NOR (conNOR) trial. The major difference between these variants of the same NOR paradigm is that stNOR includes inter-trial interval handling and removal to a holding age whereas in conNOR the rat is left to shuttle to the holding arena without any intervention from the experimenter (Ameen-Ali et al., 2012; Chan et al., 2018). Both NOR variants share initial handling into the apparatus for the acquisition phase with exploration of two identical novel objects. Thus, we were able to behaviourally dissociate the effects of distraction on memory persistence in the scPCP rat by using two variants of the same basic NOR paradigm. Our next step was to determine whether the intact memory performance of scPCP rats after the first conNOR trial would continue across the remaining 10 continuous trials of the session. The data showed clearly that whilst both groups explored objects for similar durations and were able to detect novelty above chance on each trial, scPCP performance declined significantly relative to controls from trial 3 onwards. This pattern of decreasing performance in the face of increasing, modality-specific memory from prior trials strongly supports a proactive interference effect. Thus, the continuous accumulation of object memory in the scPCP group over trials was interfering strongly with their instantaneous judgement of object novelty. Of note, the same effect was seen but to a much-reduced extent in the vehicle group. Whilst we used dietary restriction to promote conNOR performance (Chan et al., 2018), in practice many pellets were either not collected or left unconsumed by rats during the task, so we feel performance would be equally good in future studies without prior food restriction. Overall, these observations strongly support the presence of two separable deficits of cognitive control in the scPCP rat; distraction and proactive interference. The importance of this result is that two core cognitive symptoms of schizophrenia can be behaviourally dissociated in the scPCP CIAS model using two variants of the same simple, high-throughput NOR procedure.

It is worth considering the neural mechanisms that might underpin the present cognitive deficits in the scPCP model. There is substantial evidence that damage to PFC produces increased sensitivity to distraction and proactive interference. This region shows clear changes in schizophrenic patients with a strong parallel in functionally equivalent regions of frontal cortex in the scPCP model. Whilst there is good evidence for PFC dysfunction in facilitating interference there is also evidence that regions such as the perirhinal cortex (PRC) are important in this respect, particularly for object memory. Substantial evidence shows that PRC is vital for judgments of object familiarity. Data from human fMRI also show increased PRC activity for object memory retrieval under conditions of object but not spatial proactive interference (Watson and Lee, 2013) and this role for PRC in reducing object interference may be part of a wider medial temporal lobe network that includes lateral entorhinal cortex and hippocampus (Reagh and Yassa, 2014) that, in turn, communicates with PFC. These temporal lobe regions show decreased volume in schizophrenic patients and adolescents with a high risk of the disease (Roalf et al., 2017; Turetsky et al., 2003; Sim et al., 2006). Perirhinal lesions in wild-type rats produce a very similar pattern of proactive interference interfering with object memory over sequential trials in the bow-tie maze (Aggleton et al., 2010; Albasser et al., 2015) to that seen here in the conNOR performance of scPCP rats. In addition, a recent brain volume analysis in scPCP rats showed that whilst there was a general shrinkage across all regions this was particularly strong for perirhinal cortex (Doostdar et al., 2019). Thus, the strong effect of proactive interference in conNOR described here for scPCP rats may be a particular function of PRC damage in the model, which would be consistent with lesion data and results from schizophrenic patients.

In summary, the present study has shown a behavioural dissociation between effects of distraction and proactive interference on object memory by employing two variants of the same NOR paradigm in the scPCP model for CIAS. This provides a new set of high-throughput tasks with which to probe fundamental cognitive symptoms of schizophrenia in preclinical models.

## Supplementary Figure legend

**S-Figure 1.**
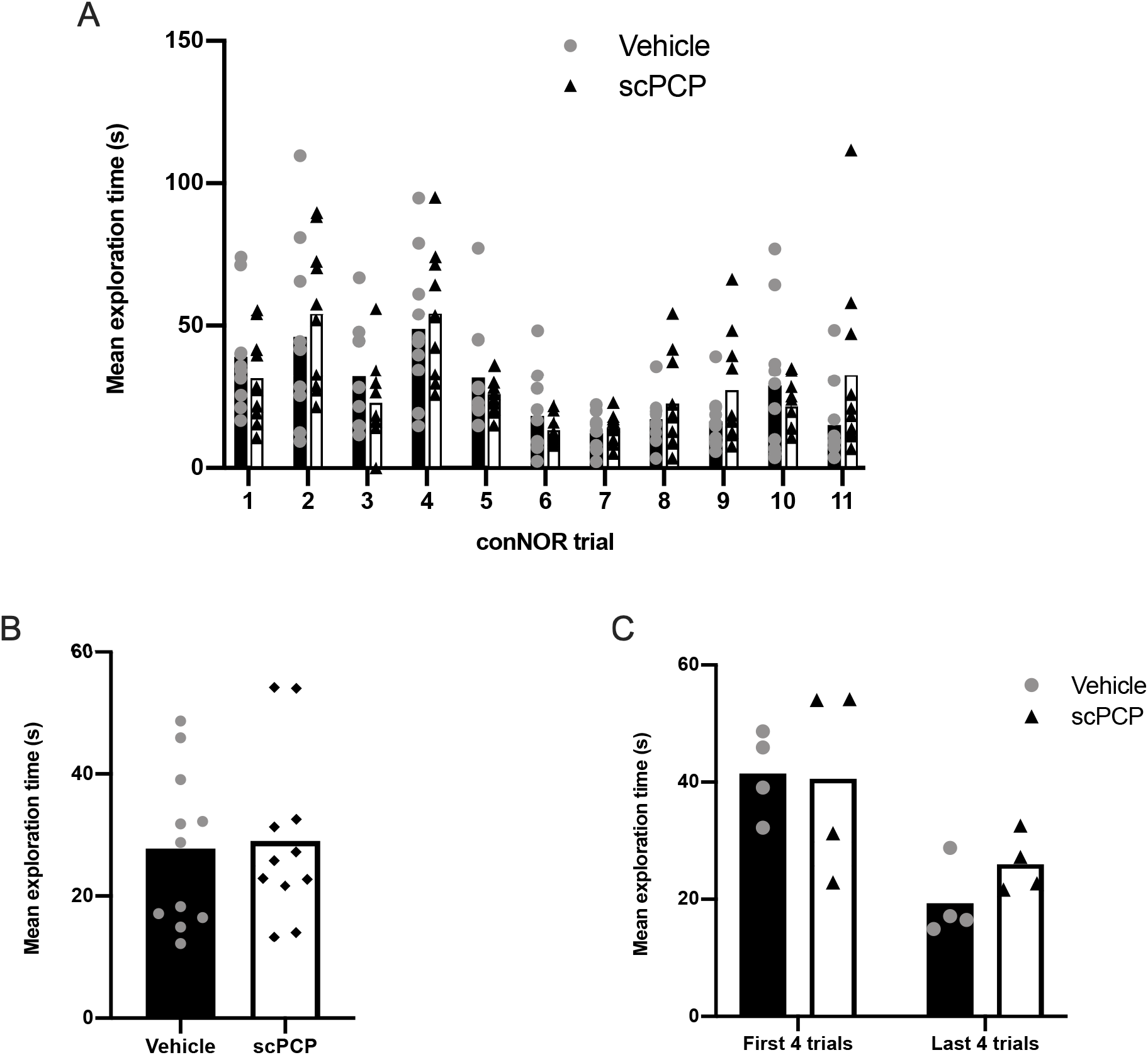
Total object exploration duration in conNOR is similar for vehicle and scPCP groups. A. Total object exploration for all conNOR trials. Exploration decreased significantly over trials but there was no effect of treatment or interaction. B. Mean exploration between vehicle and scPCP groups was similar. C. Mean exploration decreased between the first and last blocks of 4 trials but there was no effect of treatment and no interaction. N=10 for both vehicle and scPCP groups with their data presented as individual values plotted over group mean.

